# Inline SAXS-coupled chromatography under extreme hydrostatic pressure

**DOI:** 10.1101/2022.08.15.503920

**Authors:** Robert Miller, Cody Cummings, Qingqiu Huang, Nozomi Ando, Richard E. Gillilan

## Abstract

As continuing discoveries highlight the surprising abundance and resilience of deep ocean and subsurface microbial life, the effects of extreme hydrostatic pressure on biological structure and function have attracted renewed interest. Biological small angle X-ray scattering (BioSAXS) is a widely used method of obtaining structural information from biomolecules in solution under a wide range of solution conditions. Due to its ability to reduce radiation damage, remove aggregates, and separate monodisperse components from complex mixtures, size-exclusion chromatography coupled SAXS (SEC-SAXS) is now the dominant form of BioSAXS at many synchrotron beamlines. While BioSAXS can currently be performed with some difficulty under pressure with non-flowing samples, it has not been clear how, or even if, continuously flowing SEC-SAXS, with its fragile media-packed columns, might work in an extreme high-pressure environment. Here we show, for the first time, that reproducible chromatographic separations coupled directly to high-pressure BioSAXS can be achieved at pressures up to at least 100 MPa and that pressure-induced changes in folding and oligomeric state and other properties can be observed. The apparatus described here functions at a range of temperatures (0° C - 50° C), expanding opportunities for understanding biomolecular rules of life in deep ocean and subsurface environments.

## 1 Introduction

Since the early realization that hydrostatic pressure has effects on biological molecules^1^, pressure has become a useful, though not always easily accessible tool for gaining insight into basic biophysical processes^2^. In addition to being a tool for understanding phenomena such as enzymatic action, folding, and association, it is now appreciated that pressure itself is a biologically significant variable. An extraordinary portion of the biomass of our planet resides deep in the oceans, below the seafloor, and in the continental subsurface^3^. Structural biology and biophysics of these organisms should clearly be understood in the context of high pressure, yet biomolecular structural information under pressure is scarce to non-existent. As interest in deep life biology continues to grow, structural biology studies conducted under high pressure are becoming increasingly important and new instrumentation is needed to make popular biophysical techniques accessible in that regime^4^.

As an easy-to-use and versatile technique, biological small angle x-ray scattering (BioSAXS) has gained a solid foothold in the world of structural biology. A major factor contributing to its popularity is that it can be applied to a very wide range of sample conditions including high hydrostatic pressure. While high-pressure BioSAXS has been performed for some time, it has thus far remained confined to non-flowing “batch” samples^5–7^. Most contemporary SAXS facilities working at ambient pressure utilize flow cells to help mitigate the confounding effects of radiation damage. The straightforward, but very high-impact introduction of inline size-exclusion coupled to SAXS (SEC-SAXS) also allowed researchers to deal effectively with the ever-present problem of mixtures and aggregation. Moreover, SEC-SAXS provides ideal background subtractions as the elution buffer is matched precisely to the sample in the same flow cell. The importance of using SEC-SAXS to eliminate insidious problems in data interpretation has been well demonstrated in the literature consequently, the technique is available at most all synchrotron-based facilities and may well be the dominant form of BioSAXS used today^8,9^. Until the present work, high-pressure SEC-SAXS (HP-SEC-SAXS) has not been attempted.

Fortuitously, the field of chromatography has been developing high-pressure hardware in an effort to utilize smaller stationary phase particles to achieve better, faster separations^10^. Pressure gradients in columns caused by viscous flow alone can easily exceed hardware capabilities of conventional high-performance liquid chromatography (HPLC) when micron-sized particles are involved. Consequently, ultra high *performance* liquid chromatography (UHPLC) hardware has emerged on the commercial market capable of sustaining pressure gradients up to 130 MPa pressures, but only as far as the column.

For the study of the pressure effects themselves, the experimental requirements are very different from those of UHPLC: the pressure must be uniform and controllable all the way to the X-ray beam. Existing components downstream of the column are not currently designed to withstand high pressure. Consequently, there are three main technical challenges to performing ultra high *pressure* chromatography coupled to SAXS: survival of uniformly-packed column media under intense pressure cycles, X-ray transparent high-pressure flow cell design, and active high backpressure regulation.

Here, we demonstrate that size-exclusion chromatography with inline HP-SAXS measurement of biomolecules can be performed successfully at pressures of up to at least 100 MPa (1000 bar, 14,503 psi) with temperatures ranging from 4°C-50°C. Basic column performance and stability through multiple high-pressure cycles are assessed. Columns are characterized first with a set of stable protein standards showing little or no pressure effects. To study the behavior of pressure-sensitive proteins during separation, three model systems are examined: a small monomeric protein that unfolds (pp32), and two different types of cold-dissociating complexes (enolase and L-lactate dehydrogenase). We demonstrate that HP-SEC-SAXS can yield structural information in all three cases.

## 2 Results

Custom-packed chromatography columns are designated here by their packing material type (“SD” = silica diol), their pore size (120 nm or 300 nm), the column internal diameter/length in mm (5/150 or 5/300), and the packing trial number. Two column types were prepared for this study: SD120 5/150 and SD300 5/300. Column packing protocol and characterization details are given in the Methods Section (4.2) of this paper and in **Table I**. Throughout this paper, we refer to a plot of total X-ray scattering intensity as a function of elution volume or image frame number as an *elution profile*. Likewise, X-ray scattering intensity as a function of momentum transfer *q* = 4π sin θ/λ (where 2*θ* = scattering angle and *λ* = wavelength) is referred to as a *scattering profile*.

**Table 1.**
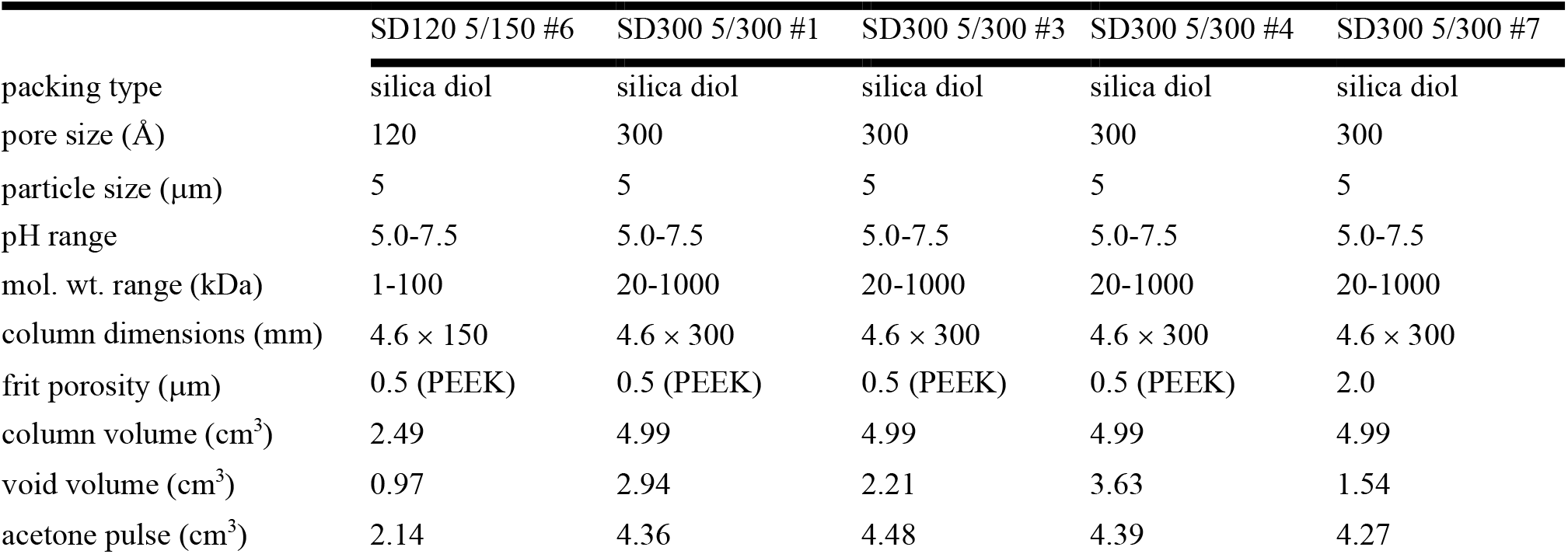
High pressure size exclusion chromatography columns

While we have attempted to minimize the pressure drop, ΔP, across the columns using standard-size chromatography particles and slow buffer flow rates, some pressure drop is unavoidable. If *P*_*inlet*_ is the pressure reported by the pump and *P*_*BPR*_ is the (reference) pressure at the outlet maintained by the backpressure regulator (BPR), then ΔP = *P*_*inlet*_ - *P*_*BTR*._ The pressure at which an observed structural change in a sample takes place happens somewhere in the pressure range *P*_*inlet*_ *to P*_*BTR*_, with the final X-ray measurement being taken at *P*_*BTR*._

Glucose isomerase, a well-studied tetrameric protein standard for SAXS, is structurally stable at pressures up to at least 300 MPa^11^. **Figure 1A** compares elution profiles of a 100 µl injection of concentrated glucose isomerase (17.9 mg/ml) at room pressure and at 102.3 MPa (*P*_*inlet*_). Buffer containing 25 mM HEPES pH 7.0, 150 mM NaCl, and 3% v/v glycerol at 23° C was used at a flow rate of 0.15 ml/min with ΔP = 8.5 MPa on column SD300 5/300 #1. X-ray exposure was continuous with each detector image collecting 2 s of data. The 13.95 keV beam was attenuated to a flux of 8 × 10^11^ ph/s to minimize radiation damage (based on lysozyme exposures at the selected flow rate). A slow leak detected during the 100 MPa measurement resulted in an effective flow rate of 0.12 ml/min, nonetheless the elution profile superimposes well with the ambient result when scaled by 1.3 to compensate for pressure-induced contrast change^11^. The mean radius of gyration of the ambient run is 34.1 ± 0.7 Å while the mean radius of gyration for the 100 MPa sample is 34.4 ± 1.0 Å. The Guinier plots of peak scattering profiles taken from these runs show no systematic deviation from linearity (**Figure S1**). Comparison of Kratky plots and difference between the two scattering profiles shows that the profiles are the same to within the noise level of this experiment (**Figure 1B,C**).

**Figure 1.**
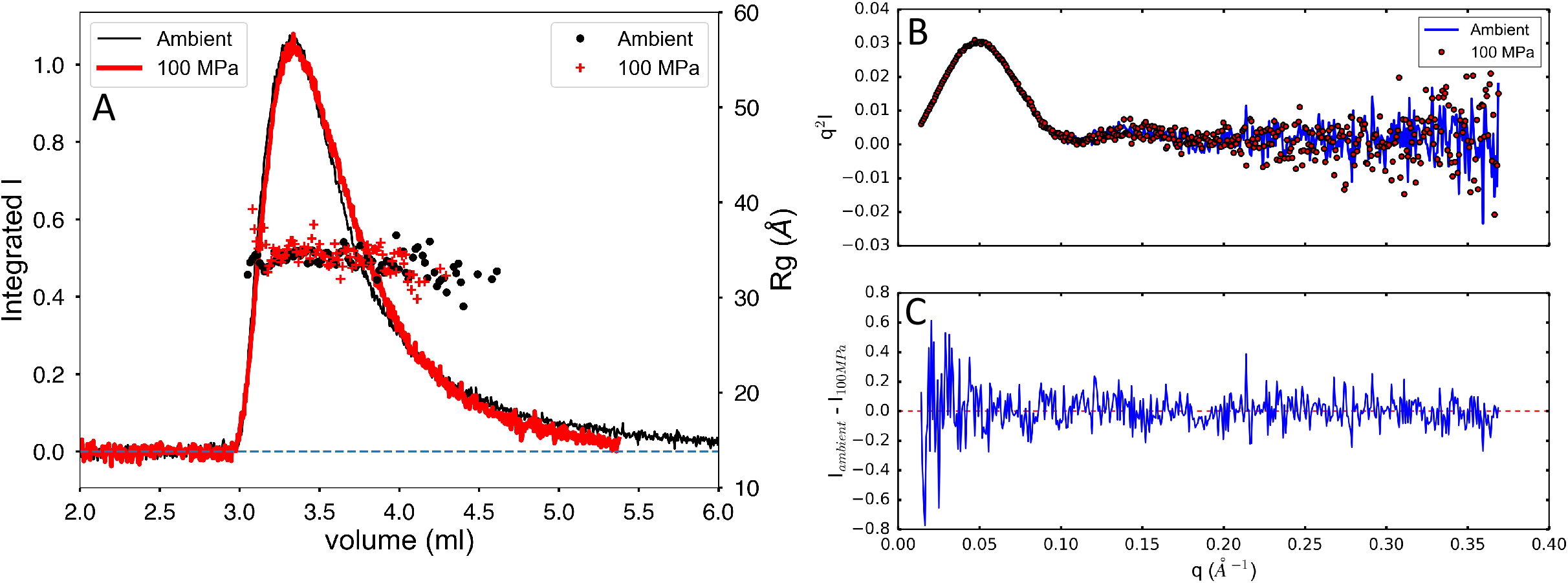
Elution profile of glucose isomerase (A) at ambient pressure and at 102.3 MPa (inlet). Solid lines are integrated intensity of the scattering profile (arbitrary units). The symbols (right-hand axis) are radii of gyration calculated by the Guinier approximation at each volume (time) point. Kratky and difference plots (B) for glucose isomerase at ambient and pressure show no significant changes with pressure.

Bovine serum albumin (BSA) is well-known as a resolvable mixture of different stable oligomeric states that can serve as a convenient standard for evaluating SEC-SAXS setups and protocol^12,13^. BSA at 20 mg/ml was run on column SD300 5/300 #3 using 25 mM HEPES at pH 7.0, 150 mM NaCl, and 1% v/v glycerol. At a flow rate of 0.15 ml/min,, ΔP = 2.4 MPa (23°C). Runs were done in the following order: 0 MPa, 100 MPa, 0 MPa “ambient final”, 50 MPa, 100 MPa (2), 100 MPa (3). The elution profiles, scaled so that the main monomer peaks overlap, are superimposed in **Figure 2A**. Column SD300 5/300 #3 resolves BSA monomer from dimer, but the very weak peak for higher oligomers sometimes visible in high-resolution separations is not resolved. The initial and final ambient results overlay closely showing that the column bed resolution did not degrade during the first pressure cycle. Though the peaks show some variability in amplitude, it is difficult to confirm any systematic change in the elution profile with pressure for this standard protein. Radius of gyration through the monomer peak shows a very slight dip that may indicate some repulsive-type concentration effects but is otherwise flat across the peaks (**Figure 2A**). Though small, the drop in radius of gyration with pressure is distinct (27.0 ± 0.04 Å ambient, 26.4 ± 0.2 Å 100 MPa) at 23°C.

**Figure 2.**
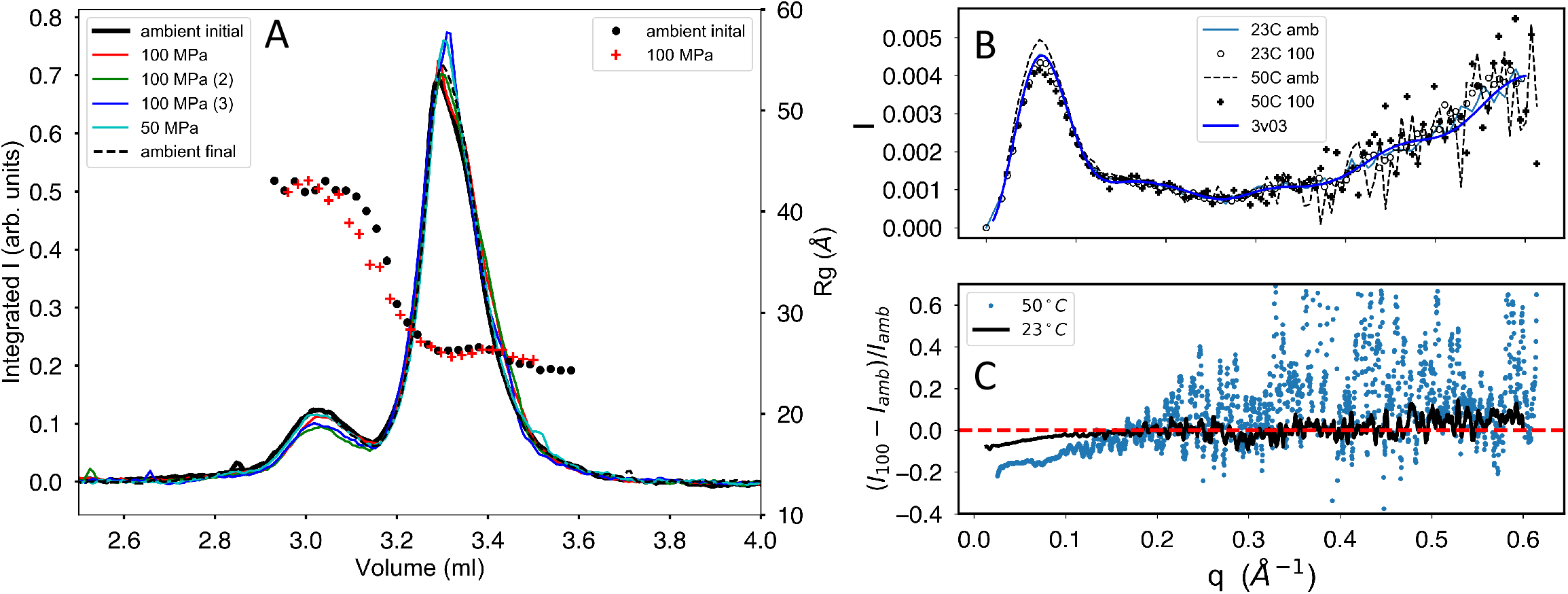
Elution profiles for bovine serum albumin through a series of sample injections at various pressures (listed in the left-hand legend box in order). (A) Monomer (main peak) is resolved under pressure from dimer with little change in the overall elution profile shape. The initial (solid black line) and final (dashed line) ambient pressure profiles overlay closely, demonstrating column bed stability through the pressure cycle. Radius of gyration R_g_ calculated along the profiles (symbols) shows plateaus at both peaks with a subtle drop in value under pressure. (B) Kratky plots show that changes in X-ray scattering with pressure are confined to the small-angle portion of the scattering profiles. (C) The change in scattering with pressure is most clearly seen by plotting relative deviation at fixed temperature.

BSA was also separated on column SD300 5/300 #7 at 13.3 mg/ml in 50 mM HEPES at pH 7.5, 150 mM NaCl, and 0.5 mM TCEP at 50°C. At this temperature a flow rate of 0.15 ml/min produced, ΔP = 1.0 MPa. The radius of gyration is higher at 50°C and shows a larger decline in R_g_ with pressure: R_g_ = 29.6 ± 0.1 Å (ambient) and R_g_ = 28.2 ± 0.1 Å (100 MPa). Comparison of Kratky plots at both 23°C and 50°C (Figure 2B) reveals that pressure-induced changes in the scattering profiles are confined to the small-angle regime (q < 0.2 Å^-1^). The changes are more clearly visible when plotted as relative deviations (*I*(*q*)_100MPa_ – *I*(*q*)_ambient_)/*I*(*q*)_ambient_) (Figure 2C). Concentration effects can be pressure dependent, though such effects are likely small at this relatively low pressure^14^. Pressure-dependent radiation damage is a more likely possibility here. Though the SAXS chromatograms return to baseline and Guinier plots are reasonably flat **(Figures S2**,**S3**), the high R_g_ values at 50°C suggest the presence of increased radiation damage with temperature. At 23°C, X-ray flux for the experiment is estimated to be 3 × 10^12^ ph/s while flux at 50°C is estimated at 2 × 10^12^ ph/s. These values are significantly higher than the previous glucose isomerase measurements. Differences in buffer composition and uncertainty in dose for these experiments limit conclusive temperature comparisons, but at fixed temperature, the dose for pressurized and ambient samples is the same. Previous studies have suggested that pressure may reduce damage-induced aggregation which could account for the observed reduction in R_g_ seen here^15^.

An example of larger-scale structural change due to pressure can be seen in pp32, the 28.6 kDa leucine-rich repeat protein (L60A mutant) used in protein folding studies^16^. A 100 µl injection of pp32 L60A at 10 mg/ml was run on buffer containing 25 mM BisTRIS pH 6.8, 10 mM NaCl, 5 mM DTT at 4° C with a flow rate of 0.15 ml/min and, ΔP = 4.5 MPa on column SD300 5/300 #3. Under pressure, P_inlet_ = 99 MPa to 101 MPa. The 14.07 keV synchrotron beam was attenuated to a flux of 4 × 10^11^ ph/s. The 100 MPa elution peak, scaled to equal area with the ambient result (**Figure 3**), appears to broaden and the radius of gyration shows slight downward drift across the peak under pressure. No leaks were detected in this run. There is a distinct increase in R_g_ from 17.68 ± 0.03 Å (0 MPa) to 19.74 ± 0.05 Å (100 MPa) consistent with unfolding. Guinier plots of pp32 L60A (**Figure S4**) at both pressures show some systematic downward curvature, especially at smallest scattering angles. Though this is a single-domain protein, the Porod-based molecular weight estimate rises from 13.1 kDa at ambient pressure to 14.1 kDa at 100 MPa. Comparison of Kratkly plots (**Figure 4A**) shows a modest, but distinct increase in the wider-angle Porod region characteristic of partial unfolding. The pair-distance-distribution function P(r) (**Figure 4B**) shows more clearly that the molecule has increased in maximum diameter significantly.

**Figure 3.**
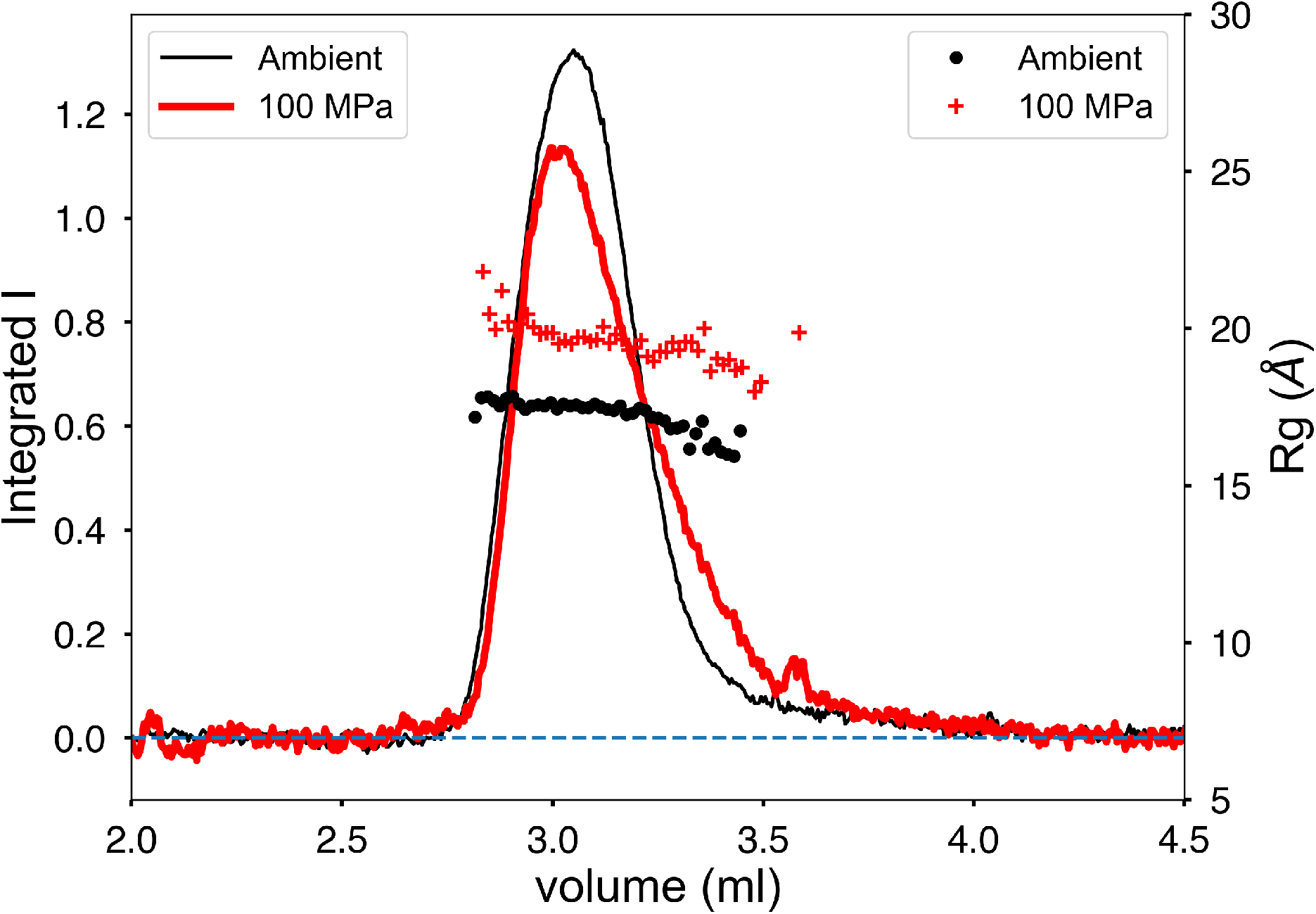
Pressure-induced unfolding of pp32 L50A during separation. The peaks have been scaled to equal area to compensate for pressure-induced contrast change. Peak broadening in the elution profile is accompanied by a significant increase in radius of gyration R_g_.

**Figure 4.**
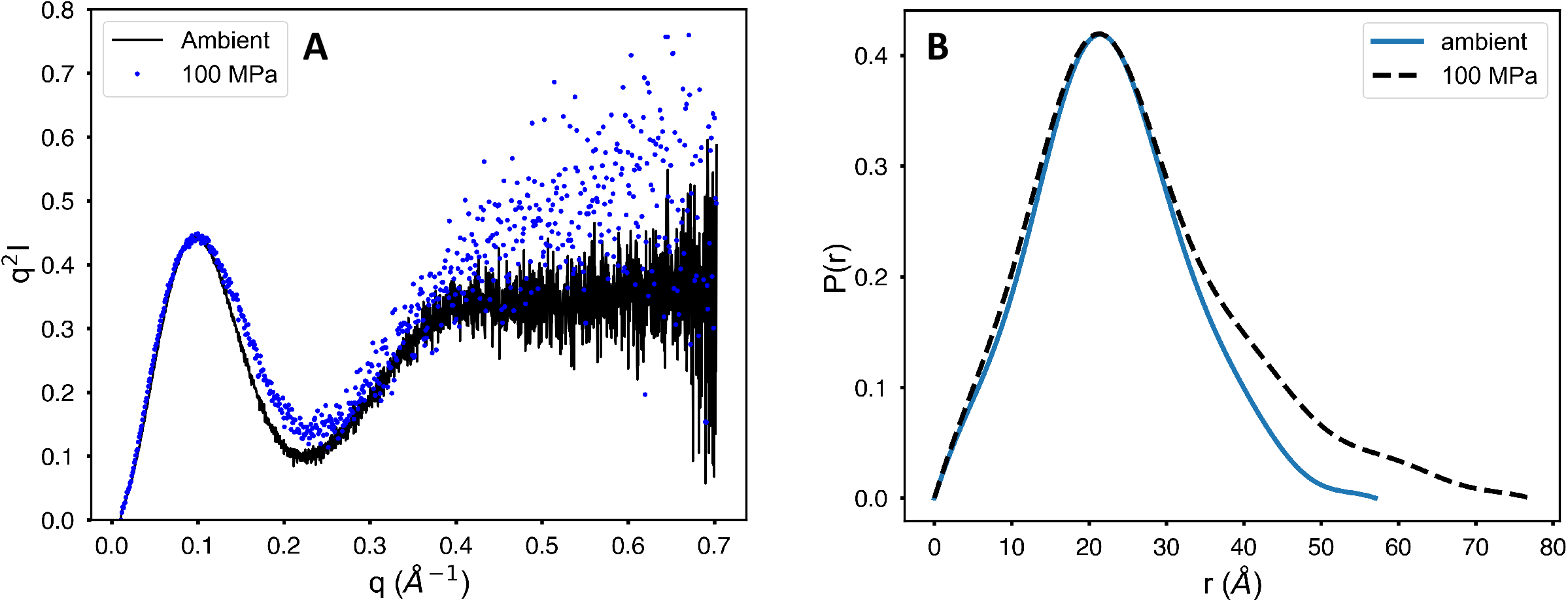
Kratky and pair distance distribution functions P(r) for pp32 in ambient and pressurized states. (A) Superimposed Kratky plots show wide-angle behavior characteristic of unfolding under pressure. (B) Pair distance distribution functions P(r) show that pressure increases the maximum diameter of the protein.

Enolase (baker’s yeast) is a 93 kDa homodimer that dissociates under modest pressure^17,18^. At 4°C and 0 MPa, enolase elutes as a single peak with R_g_ = 28.2 ± 0.1 Å and molecular weight of 85 kDa (Figure 5A). At 100 MPa, the radius of gyration drops significantly to R_g_ = 26.2 ± 0.1 Å with a drop in estimated molecular weight to 58 kDa (**Figure 5B**). The expected scattering profiles for the monomer and dimer were calculated from the crystal structures (PDB ID 3ENL) using the program FoXS^19^. The OLIGOMER program^20^ was then used to estimate the degree of dissociation at each point (frame) in the chromatogram (dots and triangles in **Figures 5A,B**). The accompanying χ^2^ plots indicate quality of fit.

**Figure 5.**
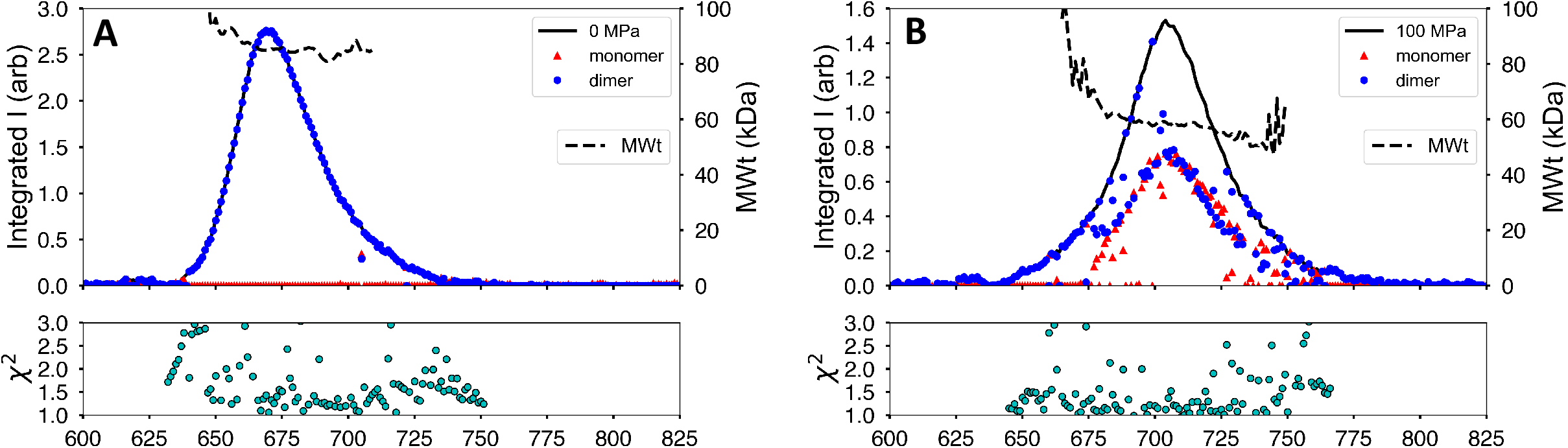
Yeast enolase becomes a mixture of oligomeric states under pressure. (A) At ambient pressure, the chromatogram is a single peak with molecular weight estimates along the elution profile that match the dimeric state (dashed line). Each scattering profile is fit to a combination of dimer and monomer crystal structures with goodness-of-fit χ2 plotted below. Circles and triangles denote the volume fractions of dimer and monomer in the profile respectively. At ambient pressure, only dimer is present. (B) Under pressure, the molecular weight (dashed line) falls below dimer and the elution profile is seen to hide a mixture of monomer (triangles) and dimer (circles).

At ambient pressure, the single elution peak is entirely composed of dimer (blue dots in Figure 5A). At 100 MPa, the sample still elutes as a single peak, but has become a mixture of dimer (blue dots) and monomer (red triangles). The leading edge of the peak appears to be dimer rich, while the main peak body is ∼50% monomer/dimer mixture. This behaviour is consistent with rapid dimer-monomer equilibrium.

The final model system studied here is L-lactate dehydrogenase (LDH, *Sus scrofa*), a homotetrameric protein that also dissociates under cold high-pressure conditions.^21^ At ambient pressure (4°C), LDH elutes as a broadened peak preceded by a small shoulder of aggregates or higher oligomers (**Figure 6A**). The estimated molecular weight is constant across the peak and closely matches the known molecular weight of the LDH homotetramer (149.61 kDa)^22^

**Figure 6.**
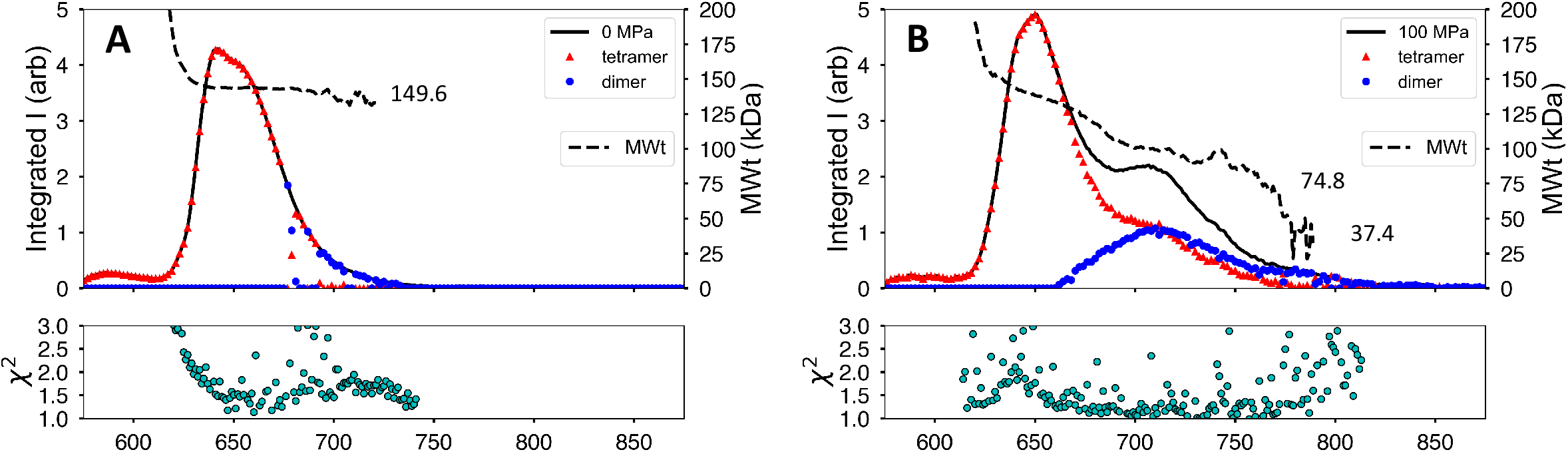
L-lactate dehydrogenase (*Sus scrofa*) chromatogram at 100 MPa shows dissociation. (A) At ambient pressure and 4° C, the protein elutes as a broad peak with molecular weight matching the tetramer (dashed line). Triangles and circles mark the computed volume fractions of tetramer and dimer respectively based on comparison with crystal structures. (B) Under 100 MPa at 4° C, the elution profile displays a second peak with lower molecular weight values. Volume fraction calculations show the second peak to be a mix of dimer and tetramer. Goodness of fit is χ2 plotted below.

At 100 MPa (4°C), an additional peak appears in the elution profile (**Figure 6B**) and the estimated molecular weight falls appreciably, though not fully to the expected dimer value (74.8 kDa). Toward the end of the second peak, the estimated molecular weight, though very noisy, settles near the monomer value (37.4 kDa). As in the previous case, the known crystal structure of porcine heart LDH (PDB ID: 6CEP)^22^ was used to compute the volume fraction of tetramer and dimer at each point along the elution profiles. LDH is commonly referred to as a dimer of dimers where the proposed solution-state dimer conserves a strong intermolecular interaction, an alpha-helical barrel between the monomeric units. The 100 MPa tetramer and dimer peaks are shown in **Figure 6B** as red triangles and blue dots respectively. By this calculation it appears that that the “dimer peak” is really a 50:50 mixture of dimer and tetramer. The late-eluting region that gives a noisy, but monomer-like molecular weight also has poor χ^2^ values in this model of only tetramer and dimer. Inclusion of a monomer model, however, did not yield improvement in that region.

## 3 Discussion

Our results show that, with care, high pressure size-exclusion columns packed in-house can show reproducible results and survive multiple pressure cycles, but must be monitored for degradation and re-packed or changed out more frequently than conventional ambient-pressure chromatography columns. Because SEC-SAXS produces well-matched buffer, it is a natural platform for investigating subtle structural changes.

The surfaces of size-exclusion particles are functionalized to resemble water with the goal of minimizing interactions between the surface and the analyte. We see no evidence of any systematic change in retention mechanism at 100 MPa in the present study. Our SD300 5/300 columns resolve monomer from dimer in BSA and, by comparison, are roughly intermediate between Superdex Increase 200 5/150 and Superdex 200 10/300 commonly used in ambient SEC-SAXS^12^.

Glucose isomerase shows no significant change at 100 MPa and can serve as a standard for testing and calibration. Bovine serum albumin (BSA) monomer is resolvable from dimer under pressure though subtle changes were observed in the small-angle part of the profile. More controlled studies are needed to access the relative sensitivity of concentration effects and radiation damage to pressure.

Protein unfolding can happen at 100 MPa, as exemplified in pp32, but such pressure sensitive proteins are probably rare. Unfolding on this scale can be seen in the Kratky plot, but is more obvious in PI, the pair distance distribution function. Spectroscopic studies of pressure-induced dissociation of protein complexes have identified two categories: complexes that reversibly dissociate and those that show hysteresis effects upon depressurization^24^. Enolase is a member of the former category since it shows rapid, reversible dissociation with little evidence of change in conformation^18^. L-lactate dehydrogenase, on the other hand, is a member of the latter category since it shows well-documented hysteresis with pressure that has been interpreted as conformational shift preventing rapid reassociation upon depressurization^21^. The differences in these two examples are very clear in HP-SEC-SAXS: upon pressurization, enolase becomes a mixture of dimer and monomer, but it remains a single peak in the chromatogram. LDH, when subjected to the same pressure, separates into two peaks representing tetramer and dimer. At the resolution used in this study we see no evidence in the SAXS profiles of conformational change in the LDH dimer.

With appropriate back-pressure regulation and sample-cell design, size-exclusion chromatography coupled SAXS can be performed repeatedly at hydrostatic pressures as high as 100 MPa while preserving column separation quality. The well-established advantages of SEC-SAXS are thus fully available for studies of biomolecules relevant to deep life and high-pressure biophysics.

## 4 Materials and methods

### 4.1 Pump and sample injection

High pressure buffer flow was supplied using a Shimadzu Nexera LC-30AD chromatography pump. Pressurized tubing was stainless steel (OD 1.58 mm, ID 0.25 mm) with a pressure rating of 138 MPa (IDEX Health & Science, LLC, Oak Harbor, WA) (**Figure 7**). For hardware and safety protection in the event failure of the back-pressure regulator diaphragm, a rupture disc was employed in the buffer flow path (RD20000, HiP High Pressure Equipment, Erie PA). Injection was accomplished using an MX Series II 2-position/6-port Ultralife™ switching valve with maximum pressure rating 103 MPa (IDEX Health & Science, LLC, Oak Harbor, WA). The valve was configured as a standard chromatography system with a 100 µl sample loop that could be loaded at room pressure using a syringe then switched inline without interrupting column flow. To maximize reproducibility during injection, the sample loop was first emptied by filling with air, then filled to maximum loop volume using an excess of sample.

**Figure 7.**
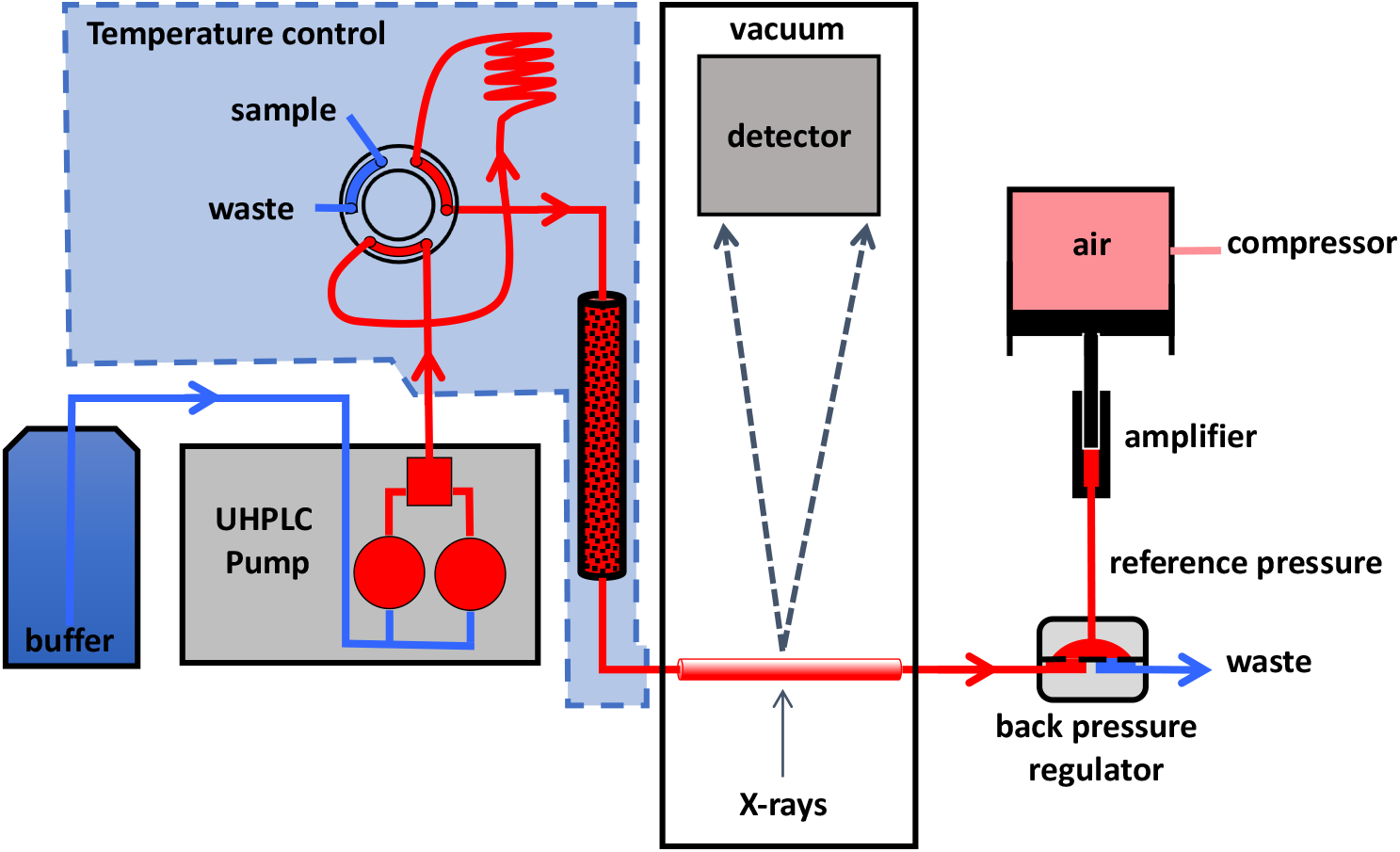
High-pressure chromatography-coupled small-angle X-ray scattering setup. Ambient pressure buffer (blue) is pressurized (red) and pumped into a temperature-controlled chamber containing the injection valve, sample holding loop, and chromatography column. After injection, sample passes through the column into a single-crystal sapphire capillary in the detector vacuum chamber where it is exposed to X-rays. Uniform hydrostatic pressure is maintained from start to end using a dome-style back-pressure regulator controlled by a pressure amplifier.

### 4.2 Column packing and characterization

The experiments reported here use diol-functionalized silica-based particles (Diol 12 nm S-5um DL12S05, Lot No. 8778; Diol 30 nm S-5um DL30S05, Ser. No. 122EA90057; YMC Co. Ltd,

Kyoto, Japan). Column hardware is PEEK (polyether ether ketone)-lined steel (PLS) (BioComp System 4.6, 15 cm and 30 cm, with 0.5 µm PEEK frits, IDEX Health & Science, LLC, Middleboro, MA) with a maximum pressure rating of 138 MPa (20,000 psi). The column hardware used for the 50°C separation was Modular Systems 4.6 mm ID 30 cm with a 2 µm frit (IDEX Health & Science, LLC, Middleboro, MA).

A 20 ml high-pressure reservoir (Teledyne SSI, State College, PA) with a short 4.6 mm ID “pre-column” was attached to the 15 cm column and filled to capacity with slurry containing an excess of packing material (∼3.5 g per 20 ml). The packing hardware used in our setup limited backpressures to below 50 MPa. Early columns loaded at APS BioCAT, in particular SD120 5/150 #3, used 20% ethanol slurry solvent with a flow rate of ∼2 ml/min resulting in a final backpressure of 12.45 MPa. Later columns packed at CHESS used a variety of solvents, but SD120 5/150 #6 used 100% isopropanol and reached a pressure of 46 MPa. After multiple volumes of isopropanol, the column was flushed with DI water giving a final backpressure of 2.3 MPa at a flow rate 0.15 ml/min (22° C). The 30 cm column, SD300 5/300 #1, was packed using 20% ethanol in water and required twice the solid material in the same 20 ml reservoir at the same flow rate and final pressure. Column backpressure with DI water was 3.6 MPa at a flow rate of 0.15 ml/min (22° C).

Back pressures during routine operation vary depending upon temperature, buffer composition, and condition of the column bed and frits. Specific values for each run are reported. Column void volume and peak characteristics were measured using 3.4 mg/ml blue dextran (D5751-1G, lot SLBP3949V, Sigma) and 1% v/v acetone (HPLC grade, Lot 170943, Fisher Chemicals). Acetone pulse experiments were conducted on freshly packed columns at ambient pressures to serve as a means of assessing column performance and monitoring long-term degradation under multiple pressure cycles of packed columns used in this paper are listed in **Table I**.

Over the course of our earliest experiments, SD120 5/150 #6 was lightly used: 12 pressure changes, 5 different samples, over a 1-year period. A final acetone pulse measurement on SD120 5/150 #6 shows no degradation in resolution (**Figure S5**). Column SD300 5/300 #1 was put into service as part of the CHESS HP-Bio facility and consequently more heavily used: 16 samples, 31 pressure changes including one accidental over-pressurization which resulted in sudden loss of the BPR rupture disk and rapid depressurization. Comparison of initial and final acetone pulse peaks shows some loss of resolution in that column at end of run (**Figure S6**).

### 4.3 High pressure flow cell

Sample flow cells for BioSAXS are normally constructed from thin fragile low-Z materials such as glass, polyimide, or mica with the goal of minimizing parasitic X-ray scattering. Single crystal sapphire capillaries have been commercially available for some time and used successfully for modest high-pressure X-ray applications [16]. The experimental cell described here (Figure S7) utilizes a 1.524 mm OD χ 1.067 mm ID χ 100 mm single crystal sapphire capillary produced by the edge-defined film-fed growth (EFG) method (SA-22979, Saint-Gobain Crystals, Milford, NH). Based on a tensile strength of 328 MPa [17], we estimate the burst pressure to be 112 MPa. The capillary was mounted inside a vacuum chamber to eliminate additional windows and air scattering (Ideal Vacuum Products, LLC, Albuquerque, NM).

### 4.4 Backpressure regulation

Liquid chromatography systems are designed to provide constant flow or, in some cases, constant pressure, but not both simultaneously. Backpressure regulation is accomplished in this work using a commercial dome-type backpressure regulator (BPR) designed to accommodate 1.5875 mm steel chromatography tubing with sample pressures up to 138 MPa (U20L Series, Equilibar, Fletcher, NC). Because the unit functions at a 1:1 pressure ratio between sample and reference, we used hydrostatic pressure rather than gas pressure for safety reasons (**Figure 7**). Our tests found that pressure under realistic constant flow conditions of a chromatography experiment varied by 0.16% or less over a 60 min time period for pressures above 20 MPa, a level comparable to the stability of the reference source. Reference hydrostatic pressure was supplied using a commercial pressure intensifier system (HUB440 High Pressure Generator, Pressure BioSciences Inc., South Easton, MA). Unless otherwise stated, reported pressures in this paper are “pre-column” buffer pressures, *P*_*inlet*_, registered by the chromatography pump.

### 4.5 Temperature regulation

Temperature regulation was achieved with a cooling-incubator (SPX-70BIII, Faithful Instrument (Hebei) Co., Ltd, China) capable of housing the liquid chromatography columns and injection valves and maintaining a constant temperature over the range 0-60 °C (Figure S8). The UHPLC pumps are outside of the cooling-incubator; tubing leading from the UHPLC pump to the cooling-incubator is uninsulated but is allowed sufficient time within the cooling-incubator system to equilibrate before reaching the chromatography system (**Figures 7, S8 3**). Tubing leaving the cooling-incubator is insulated by a chiller-regulated water jacket. The tubing and sample cell within the vacuum chamber are unregulated but, given an average flow rate of 0.1-0.3 mL/min and an approximate total volume of 0.035 mL, the sample reaches the X-ray beam in **7** seconds.

### 4.6 Synchrotron beamline characteristics

Experiments at CHESS beamline ID7A (HP-Bio) were conducted shortly after the completion of the CHESS-U upgrade to the synchrotron ring, consequently conditions varied as the ring current and beam characteristics were gradually ramped up during the commissioning period. In all cases, a 0.25 mm χ 0.25 mm beam was used at 14 keV (0.88 Å) with flux ranging from 4 χ 10^11^ to 3 χ 10^12^ photons/s (50 ma – 100 ma positron ring current). X-ray exposure is continuous with each detector image corresponding to 2 s of exposure. Sample-to-detector distances ranged from 15–0 - 1700 mm (SAXS) with earliest measurements being collected on a Pilatus 100k-s detector (Dectris, Switzerland). Exposures were normalized in all cases using transmitted beam as measured by PIN diode in the beamstop. Final data were collected on a in-vacuum EIGER 4M detector (Dectris, Switzerland) spanning a q range of 0.01Å^-1^ to 0.7 Å^-1^.

### 4.7 Samples

Glucose isomerase used in this study was from the same batch as used in evaluating our static HP-SAXS system. The preparation protocol can be found in the paper describing that work^11^. The variant pp32 L60A was produced as a C-terminal His-tag fusion, purified according to previously described protocol^16,25^. The following proteins were purchased from MilliporeSigma (St. Louis, MO) and used without further purification: enolase from baker’s yeast (E6126, lot #SLCC98590) and L-lactate dehydrogenase from hog muscle (10107085001, lot #14100847).

### 4.8 Data processing

Detector images were reduced to scattering profiles using the BioXTAS RAW software^27^. For runs with non-drifting chromatographic baselines, buffer data were taken typically from 90 frames prior to the peak. Cases of baseline drift were linearly corrected with buffer frames before and after the peak using RAW’s Baseline Correction feature. Single averaged, buffer-subtracted profiles were generated by symmetrically choosing frames from the peak down to half-maximum.

Radius of gyration was calculated by the customary Guinier analysis as implemented in RAW. Molecular weights were calculated based on corrected Porod volumes^28^. Integration *q* cutoff was 8/*R*_*g*_ Å^-1^ with a macromolecular density assumed to be *ρ*_*m*_ = 0.00083 kDa/Å^3^. Note that in the Porod volume expression,

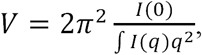

pressure-dependent pre-factors of *I*(*q*), such as contrast, cancel. Computation of molecular weight from Porod volume depends upon the average mass density of protein: *MWt* µ *ρ*_*m*_ *V* ^28^. While *ρ*_*m*_ is potentially pressure sensitive, proteins tend to be much less compressible than water (typically by a factor of 10)^29^. Since water compresses by only 4% at 100 MPa, any systematic error introduced by variation of the protein mass density with pressure will likely be well below the typical 10% error inherent in molecular weight estimates.

Pair distance distribution functions, P(r), were calculated using GNOM^30^. Calculation of volume fractions in mixtures based on known scattering components were done with OLIGOMER^31^ with *q*_*max*_ = 0.2 Å^-1^ and the constant “-cst” option invoked. Predicted profiles calculated from known PDB structures were obtained from the FoXS program^19^ except in cases where solvation parameters were being adjusted, in which case CRYSOL was used^20^. Goodness of fit measures, χ^2^, are reported as calculated by the respective programs.

SAXS profiles for monodisperse samples in this paper have been deposited in the Small Angle Scattering Biological Data Bank (SASBDB) with the following accession numbers: SASDP47, SASDP57; SASDP67, SASDP77, SASDP87, SASDP97; SASDPA7, SASDPB7.

## 5 Supplementary material description

**Figure S1.**
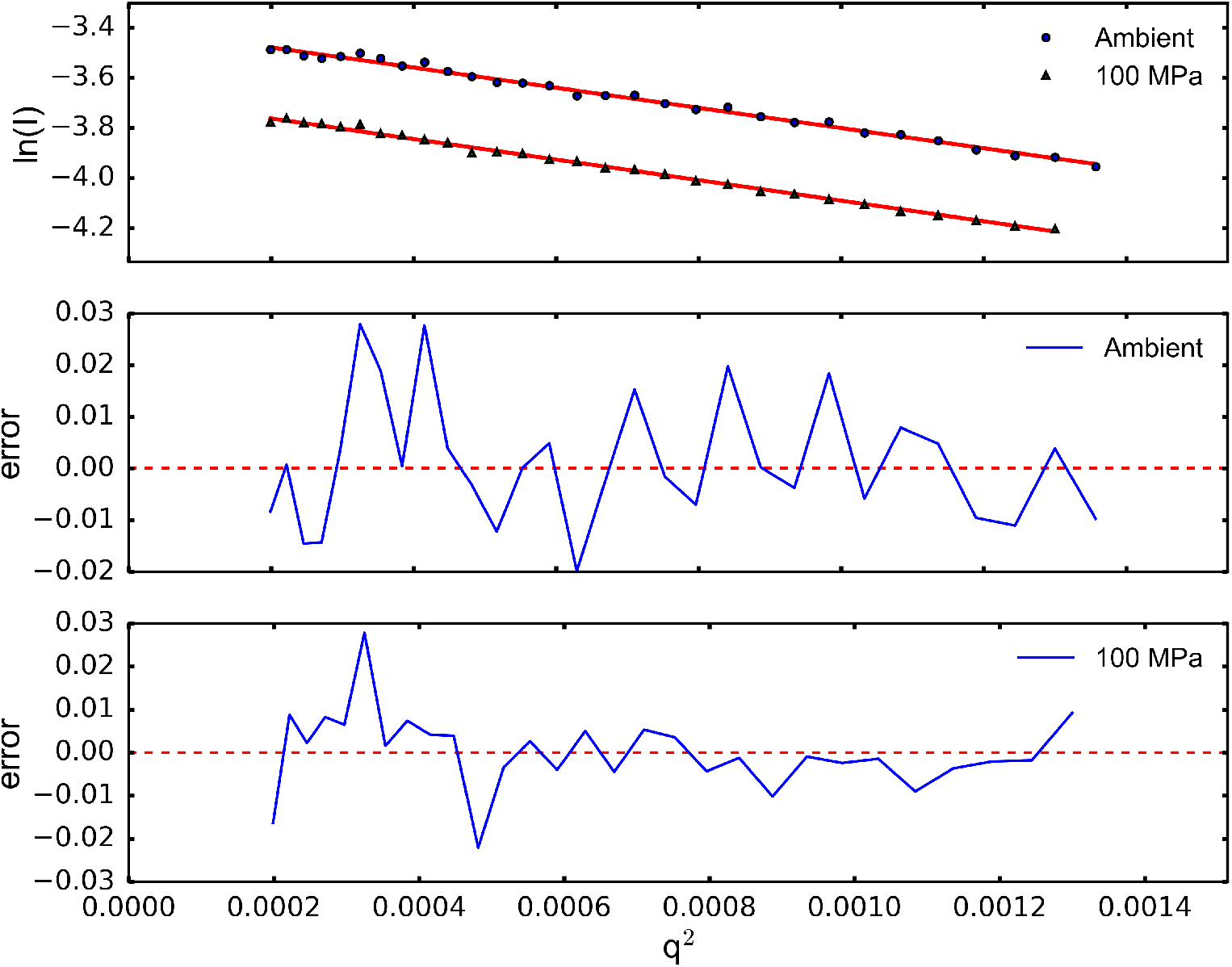
Guinier analysis of glucose isomerase X-ray scattering profiles at ambient and 100 MPa pressure. Plots of deviation from linearity (error) are in absolute units.

**Figure S2.**
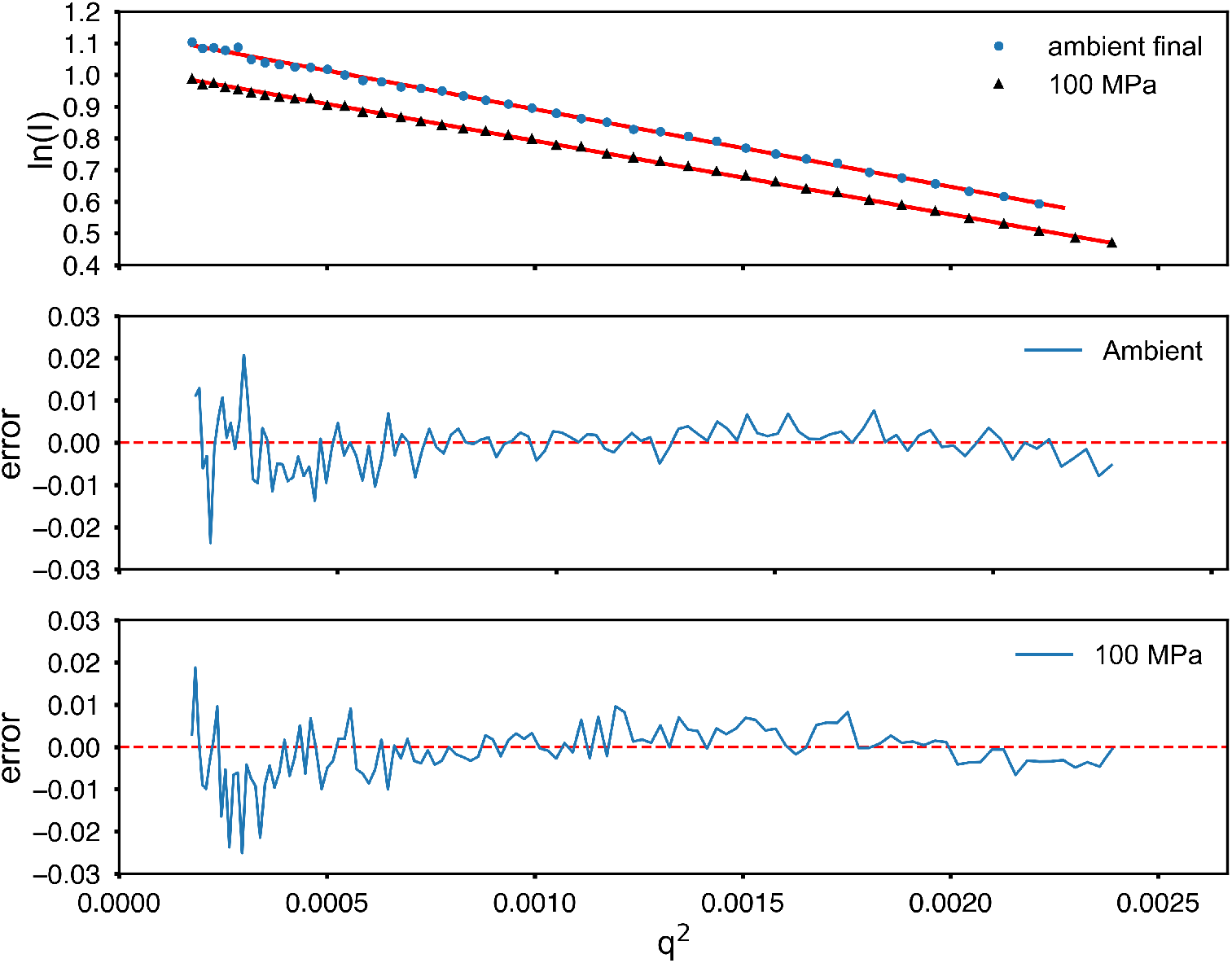
Guinier analysis of bovine serum albumin monomer X-ray scattering profiles at ambient and 100 MPa pressure, 23 °C. Plots of deviation from linearity (error) are in absolute units.

**Figure S3.**
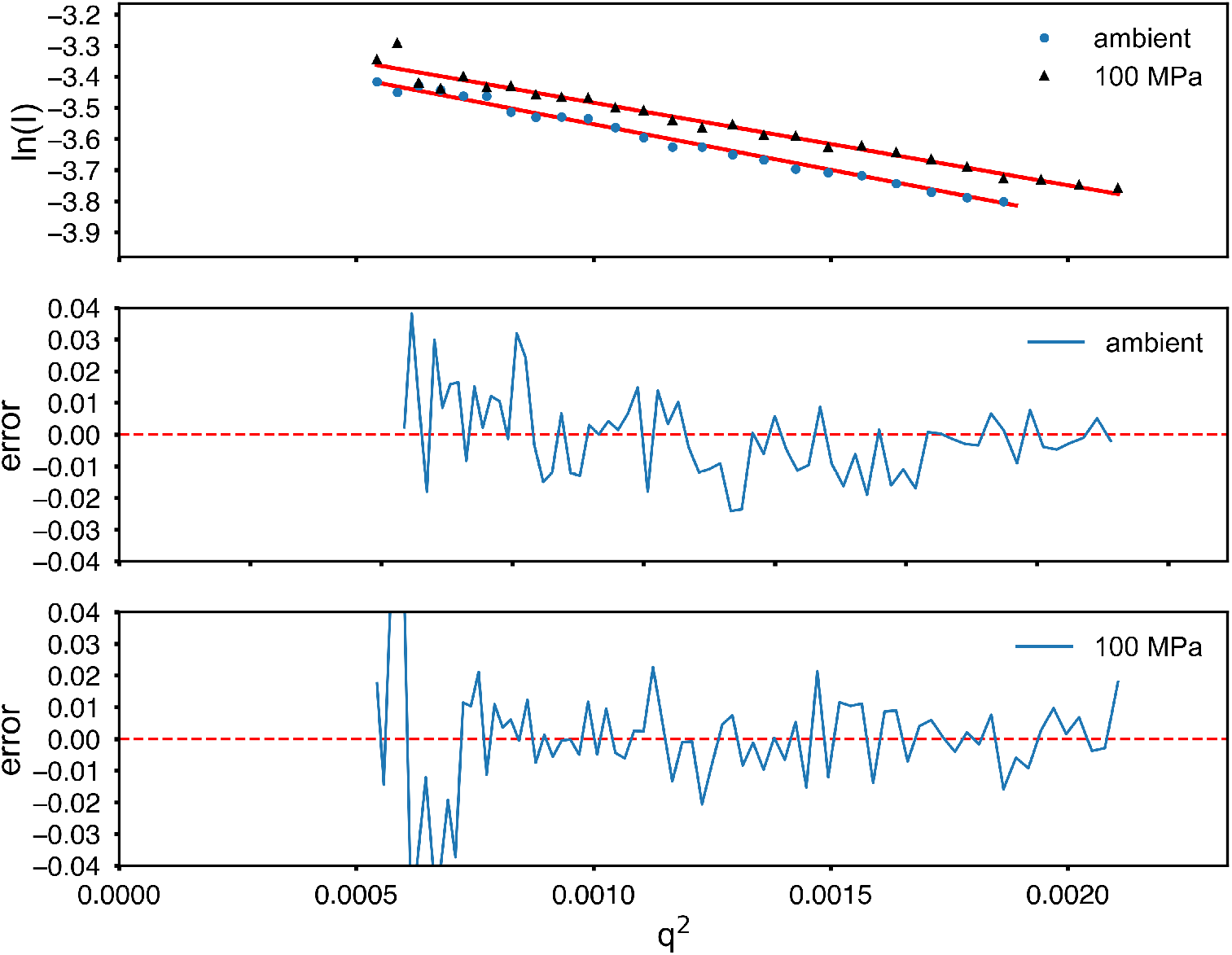
Guinier analysis of bovine serum albumin monomer X-ray scattering profiles at ambient and 100 MPa pressure, 60 °C. Plots of deviation from linearity (error) are in absolute units.

**Figure S4.**
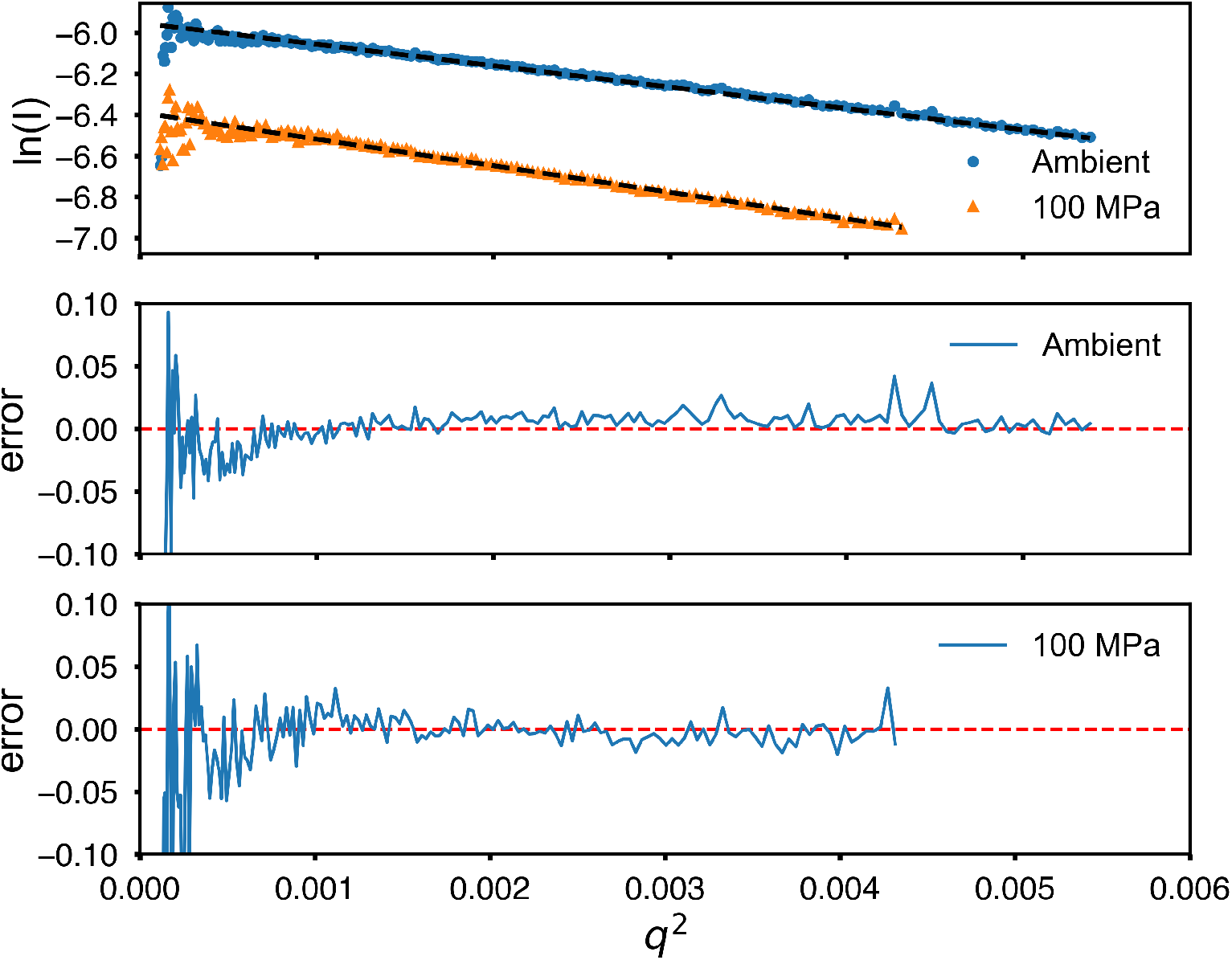
Guinier analysis of pp32 X-ray scattering profiles at ambient and 100 MPa pressure. Plots of deviation from linearity (error) are in absolute units.

**Figure S5.**
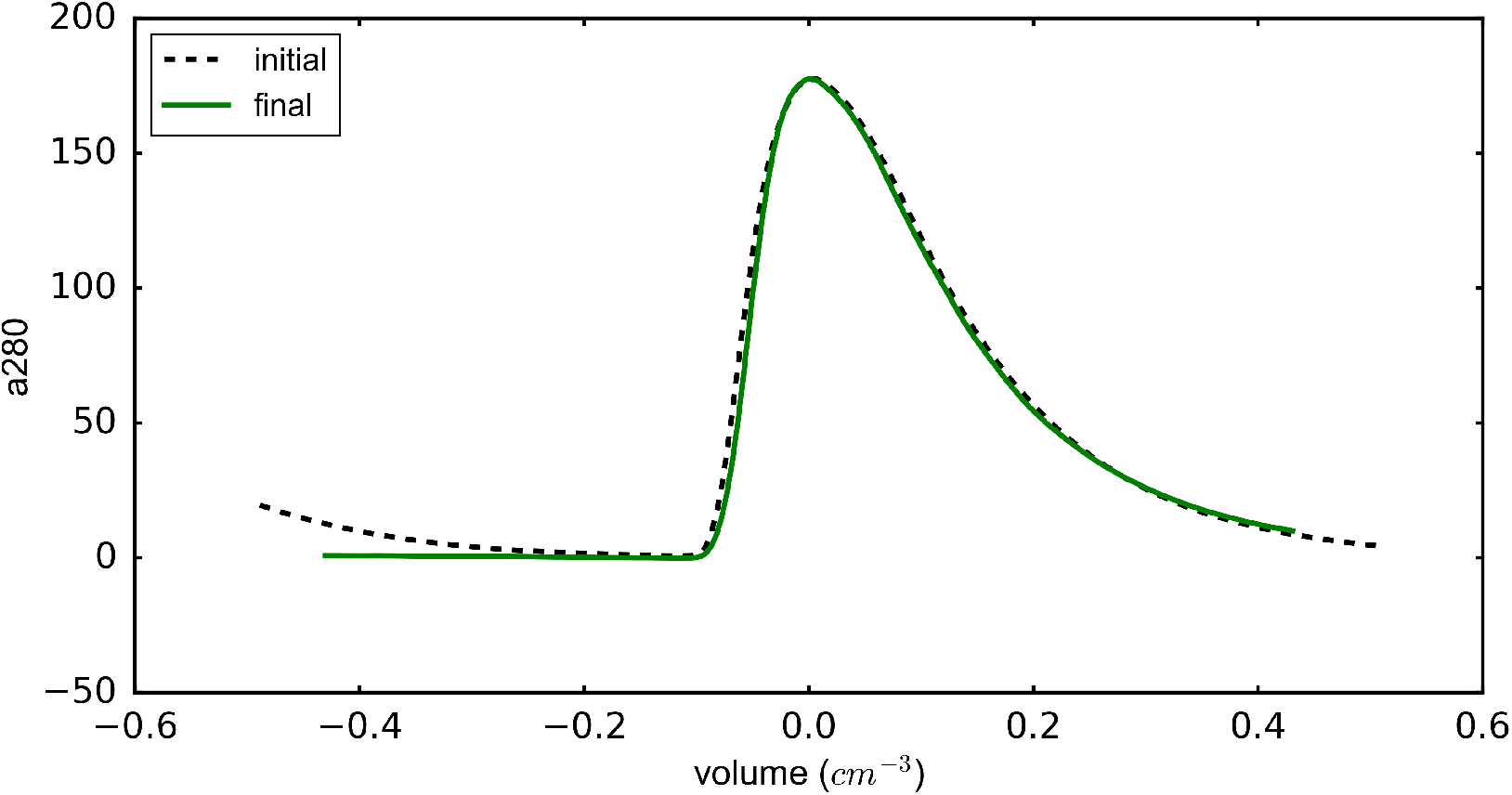
Acetone pulse reproducibility on column SD120 5/150 #6 (Table I) after light repeated use: 12 pressure changes, 5 different samples, over a 1-year period. Close superposition indicates no bed damage occurred during pressure cycles.

**Figure S6.**
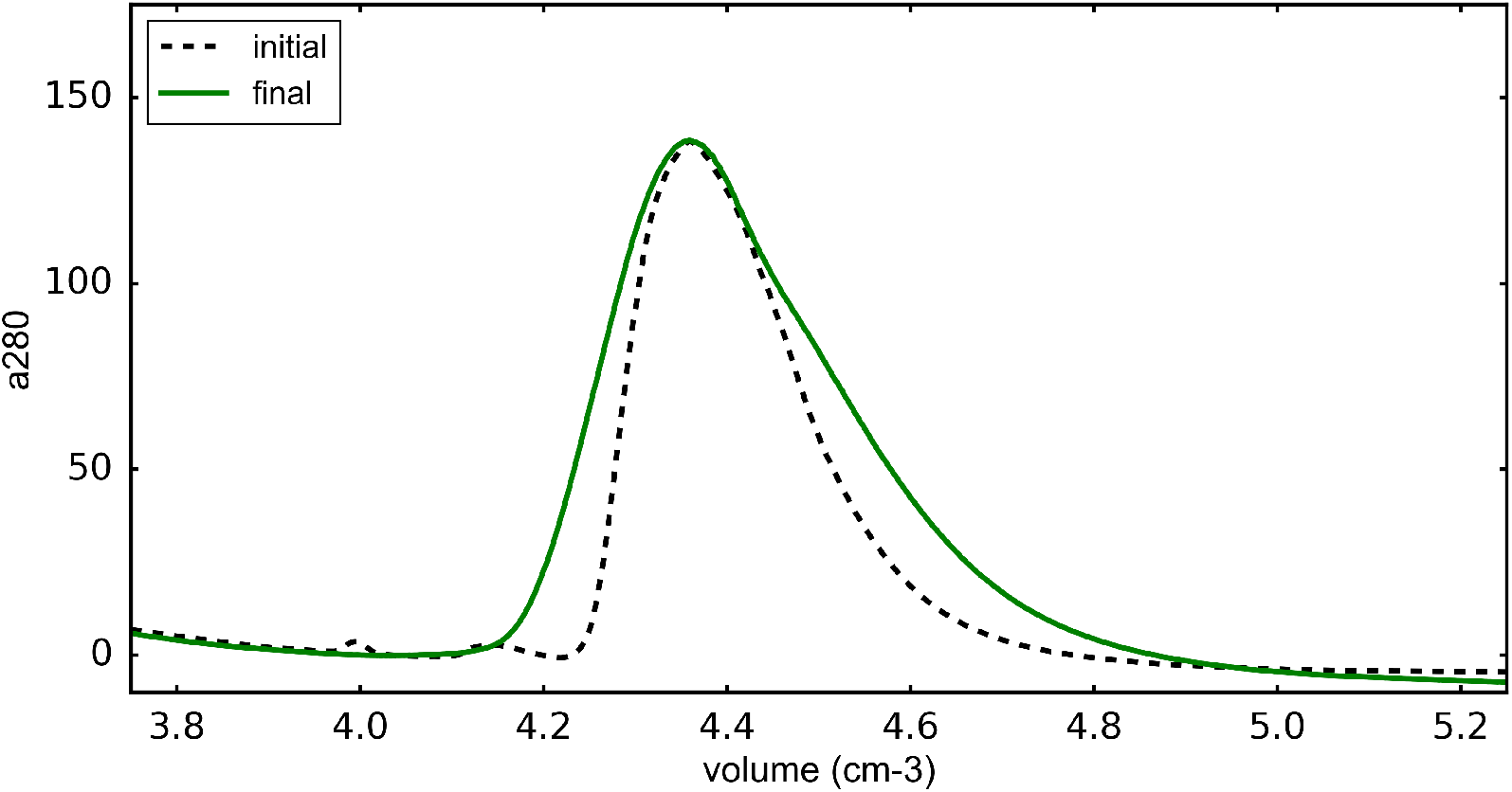
Acetone pulse reproducibility on column SD300 5/300 #1: 16 samples, 31 pressure changes including one accidental over-pressurization that resulted in sudden loss of the BPR rupture disk and rapid depressurization. Broadening of the final peak indicates some bed damage.

**Figure S7.**
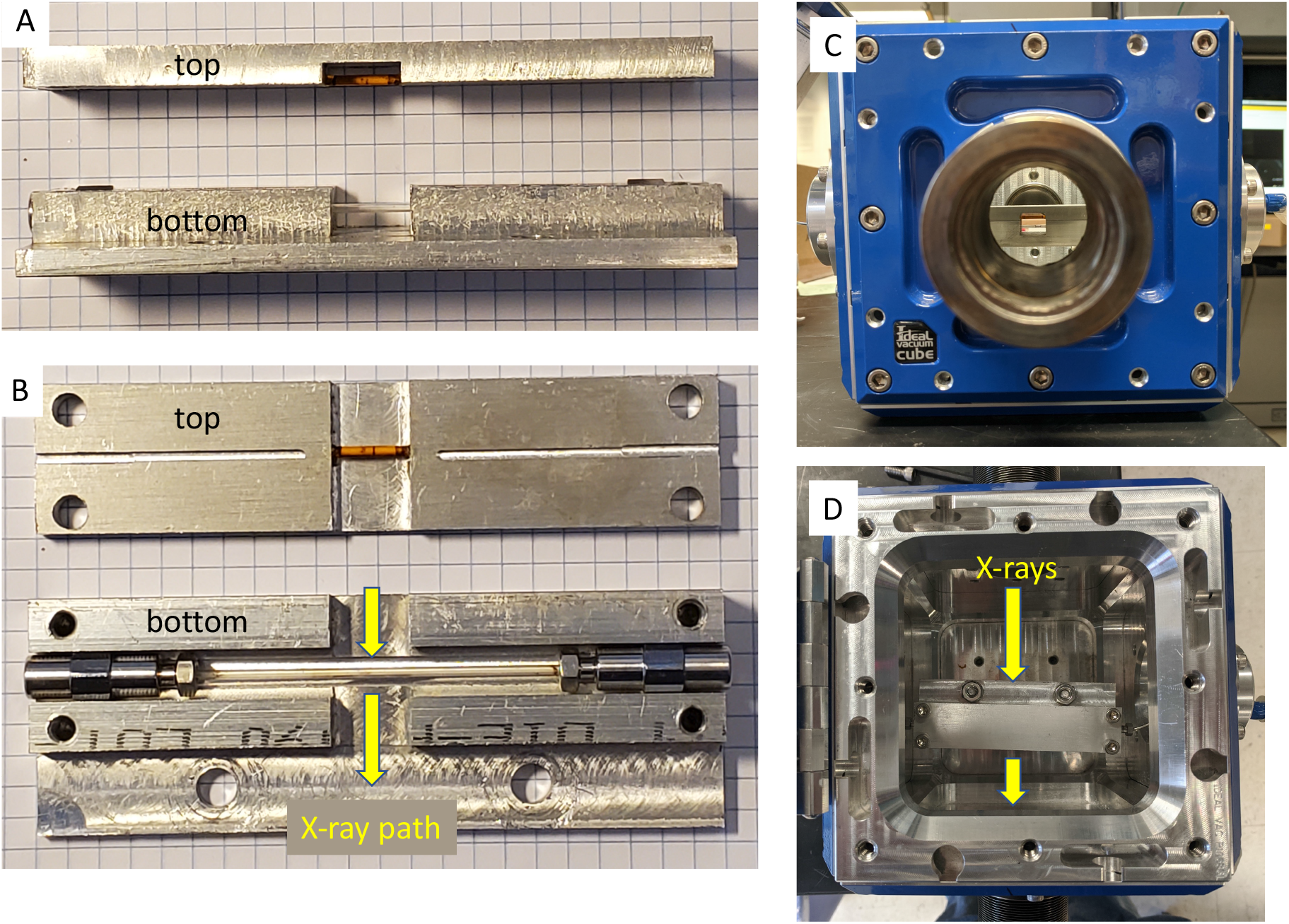
Sapphire high-pressure capillary housing. Side view (A): top cover (top) contains a small plastic capillary of silver behenate that can be lowered into the beam for calibration purposes. Bottom half (bottom) houses the single crystal sapphire capillary that connected to steel UHPLC tubing via fittings and couplers and is seated in an aluminium block in such a way that the hex faces of the couplers prevent accidental twisting during assembly (see top view B). The whole unit is secured inside a small vacuum chamber (Ideal Vacuum Products, Albuquerquem NM) that is connected to the beamline via KF 40 bellows (front view (C), top view (D)).

**Figure S8.**
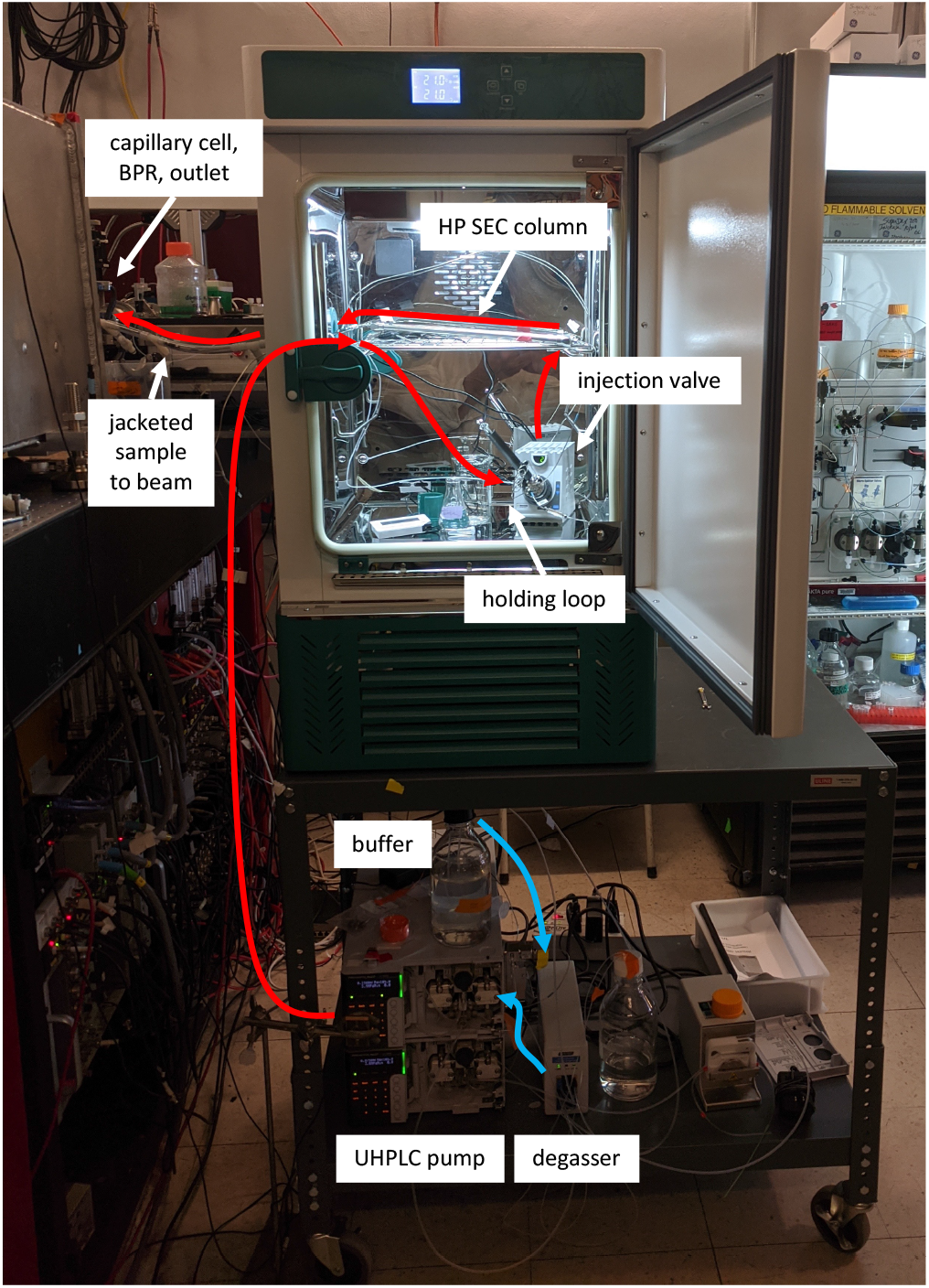
Synchrotron beamline setup. Low-pressure fluid is in blue, high-pressure fluid is in red. Buffer is drawn through a degasser (bottom) and pumped at high pressure into the temperature-controlled enclosure (top).The high-pressure injection valve inside the chamber (bottom shelf) allows ambient-pressure sample loaded in a holding loop to be pressurized and injected into the HP-SEC column. Once through the column, a chiller jacket (left) maintains the temperature of the pressurized sample until it reaches the X-ray beam enclosure (left, not shown) and exits through the back pressure regulator (left, not shown).

## Acknowledgements

Generous help and advice from Srinivas Chakravarthy and Jesse Hopkins (APS BioCAT) made the first tests of this technology possible during the CHESS-U dark period. Sriramya Nair (Cornell) and Sol Gruner (Cornell) also provided critical advice on high-pressure sample cell construction. Special thanks also to YMC-America, Inc. for generously providing sample packing material for our first experiments. We are also indebted to Equilibar Precision Pressure Control for the loan of a prototype BPR for early testing. Durgesh Rai (Cornell) provided expert advice on high-pressure equipment. Many thanks to Siwen Zhang and Cathy Royer (RPI) for providing pp32 L60A. This research used resources of the Advanced Photon Source, a U.S. Department of Energy (DOE) Office of Science User Facility operated for the DOE Office of Science by Argonne National Laboratory under Contract No. DE-AC02-06CH11357. The project was supported by grant 9 P41 GM-103622 from the National Institute of General Medical Sciences of the National Institutes of Health. This work was also supported by National Institute of General Medical Sciences Grant GM124847 to N.A. The majority of this research was conducted at the Center for High Energy X-ray Sciences (CHEXS), which is supported by the National Science Foundation under award DMR-1829070, and the Macromolecular Diffraction at CHESS (MacCHESS) facility, which is supported by award 1-P30-GM124166-01A1 from the National Institute of General Medical Sciences, National Institutes of Health, and by New York State’s Empire State Development Corporation (NYSTAR)

## Notes

### Competing Interest Statement

The authors have declared no competing interest.

